# A synthesis on alien mammals threatened in their native range

**DOI:** 10.1101/2024.03.04.582492

**Authors:** Lisa Tedeschi, Bernd Lenzner, Anna Schertler, Dino Biancolini, Carlo Rondinini, Franz Essl

## Abstract

Many alien species are safe in their native ranges; however, some are threatened, posing a challenging conundrum for conservation and invasion science. We focused on alien threatened mammals, examining their distribution, pathways, threats, and conservation strategies. We also reassessed their IUCN Red List category to evaluate the effect of including alien populations in extinction risk assessments. Of 242 alien mammals, we identified 41 aliens that are threatened, classified as critically endangered (19%), endangered (27%), or vulnerable (54%). They were primarily introduced for hunting and exchanged within Asia, with introduced ranges concentrated in eastern Australia. They were subject to multiple threats, most notably biological resource use. Including alien populations in the categorization assessments reduces extinction risk of 22% of the species. We highlighted the conservation value of alien populations for threatened mammals. However, conservation managers and extinction risk assessors shall carefully consider their use, to avoid negative impacts on native biodiversity.

## INTRODUCTION

Mankind has become the dominant force shaping Earth’s surface and its biological processes, creating an extinction crisis, with many wild populations declining and an estimated one million species facing risk of extinction globally (IPBES 2019). Humans have broken down biogeographic barriers through increased international trade and globalization, resulting in an unprecedented rise in species moved beyond dispersal barriers (Blackburn et al. 2011, Seebens et al. 2017, 2023, IPBES 2023). Species introduced by humans (alien species) play a crucial role in the biodiversity crisis, contributing to at least 60% of global extinctions (IPBES 2023). Alien species can spread and eventually threaten human livelihoods and biodiversity, thus being labelled as invasive (Blackburn et al. 2011, IPBES 2023).

Globally, 242 alien mammal species are known (Biancolini et al. 2021), and several are causing substantial impacts on the environment or human well-being (Tedeschi et al. 2022, IPBES 2023). Many alien mammals are widespread in their native range, while some may be declining and ultimately facing the risk of extinction. Previous research focused mainly on species-specific cases (Lees & Bell 2008, Garzón-Machado et al. 2012, Cassinello 2018), few species (Marchetti & Engstrom 2016, Gibson & Yong 2017) or countries (Baquero et al. 2023), while an updated global synthesis is lacking. Alien populations of mammals threatened in their native range represent a conservation conundrum, as they can be detrimental to biodiversity and ecosystem services and at the same time be important conservation assets (Marchetti & Engstrom 2016). For example, they can serve as supply of traded species, thereby reducing pressure on native populations (Gibson & Yong, 2017), or preserve landscapes (for instance, by grazing) replacing lost native species with similar ecological roles (Cassinello 2018).

Here, we provide a synthesis on alien mammals threatened in their native range, addressing the following questions: i) How many mammals threatened in their native range have established alien populations? ii) What is their native and alien geographic distribution? iii) What are the introduction pathways of those populations? iv) What are the causes of threat and which conservation measures are undertaken?; and v) How would global assessments of extinction risk change if alien populations were included?

## MATERIALS AND METHODS

For identifying alien mammals and their distribution, we used the Distribution of Alien Mammals database (DAMA; Biancolini et al. 2021), which covers 230 alien mammal species with established populations globally. We retrieved the native distributions and the extinction risk assessments from the IUCN Red List of Threatened Species (hereafter “Red List”, IUCN 2023). After resolving taxonomic differences between databases following the IUCN taxonomy (Supplementary Methods), we identified alien mammals that are threatened in their native range (henceforth, “alien threatened mammals”) as those classified as vulnerable (VU), endangered (EN), critically endangered (CR), and extinct in the wild (EW) by the Red List (Supplementary Methods). We kept the DAMA ranges for visualization purposes (Supplementary Methods). To test if the proportion of alien threatened mammals statistically differed from the proportion of alien or threatened mammals while accounting for non-independence in the observations, we used McNemar’s exact test (Supplementary Methods; McNemar 1947).

To counteract forthcoming extinction threats (e.g., fragmentation and habitat loss), some species have been introduced geographically close to the current natural range of the species, within suitable habitat and eco-geographical area (IUCN/SSC 2023, IUCN 2022a). Those are “benign introductions”, i.e., carried out for conservation purposes following IUCN Guidelines (Supplementary Methods; IUCN/SSC 2013, IUCN 2022a), and should be included in the assessment of global species extinction risks along with wild populations inside the native range. In our study, we did not consider benign introductions already included in Red List assessments. However, we considered benign introductions not included in Red List assessments or other introductions performed for conservation purposes not based on IUCN Guidelines.

We extracted pathways of introduction for each alien range polygon, as the same species could have had several alien polygons in the same continent but with different introduction pathways, from DAMA (Biancolini et al. 2021). We extracted species’ Red List category, countries of occurrence, threats, and conservation measures (Supplementary Methods; IUCN 2023). For threats and conservation measures, we used the highest hierarchical level provided by IUCN (Level 1; Supplementary Methods; IUCN 2012a, 2022b) and we aggregated all threats of lower levels to the corresponding highest level (Supplementary Methods). We assigned the continents of occurrence to each species by intersecting its native and alien countries of occurrence with continents (Supplementary Methods).

We evaluated if including all alien populations of threatened mammals would modify the outcomes of the assessments by applying the Red List assessment criteria (IUCN 2012b, 2022a). We accessed the Red List assessment (https://www.iucnredlist.org/) on 19.12.2023 and retrieved data on year of assessment, Red List category and criteria, and every other useful information. We then combined this data and the information on alien ranges (Biancolini et al. 2021) to perform Red List assessments for all study species following the IUCN guidelines (IUCN 2022a). We compared the original Red List assessments of global extinction risks (which do not always include the alien populations) and our re-assessments (based on native plus all alien populations) to illustrate differences in the resulting extinction risk category. Lastly, we used the Red List Index (RLI), which shows trends in the status of taxa, calculated based on the formula in Butchart et al. (2007) to quantify this change (Supplementary Methods). All analysis and data visualization were performed in R (R Core Team 2023). The full list of R packages used is available in the Supplementary Methods.

## RESULTS

### Number, taxonomic distribution, and threat categories of alien threatened mammals

We identified 53 alien threatened mammals (Supplementary Results), of which 12 species were solely benign introductions (IUCN/SSC 2013, IUCN 2022a) included in Red List assessments and were therefore excluded from our analyses (Supplementary Results). We thus analyzed 41 alien threatened mammals (18% of the total pool of 230 globally established alien mammals; Biancolini et al. 2021), a proportion significantly different from the proportion of alien (not threatened) or threatened (not alien) mammals (McNemar’s exact test p-value < 0.01, 95% CI 0.18–0.20). Alien threatened mammals are distributed across 10 orders, the most numerous one being Artiodactyla (n = 15 species), followed by Primates (n = 11), and Diprotodontia (n = 6) (Figure 1A, Supplementary Results). On the family level, the most important ones were Cercopithecidae (n = 7 species), followed by Bovidae and Cervidae (both n = 6) (Figure 1B, Supplementary Results). A total of 8 study species (20%) are Critically Endangered, 11 (27%) are Endangered, and 22 (53%) are Vulnerable (Figure 1); no alien threatened mammals are assigned to other Red List categories.

**Figure 1.**
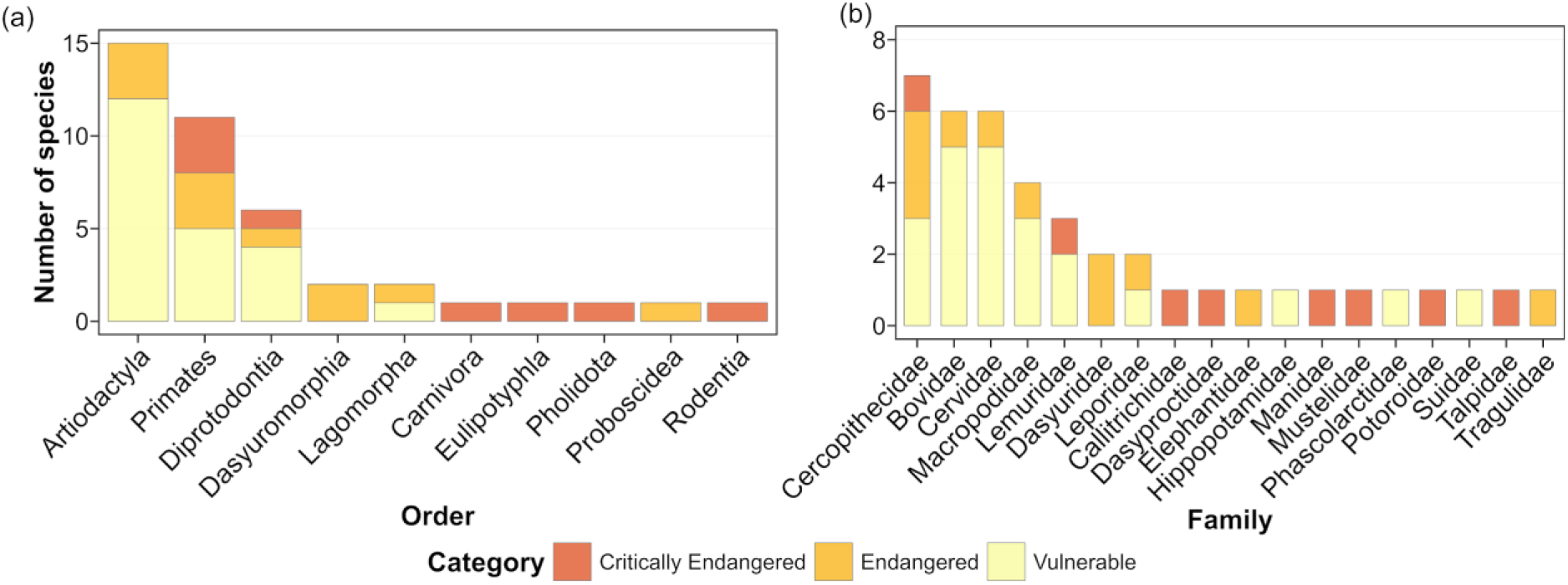
Taxonomic distribution of the 41 alien threatened mammals by (a) order and (b) family. The Red List categories of the species are shown. Red List Categories are indicated with their code: CR (Critically Endangered), EN (Endangered), and VU (Vulnerable).

### Distribution, continental flows, and introduction pathways

The distribution of the native and alien ranges of alien threatened mammals shows distinct patterns. The majority of native ranges are found in continental and insular Southeast Asia, while individual native ranges are distributed across substantial parts of all other continents except for South America (Figure 2A). Alien ranges are mainly found in eastern Australia and insular Southeast Asia, with individual alien ranges being restricted to Europe, and to small fractions of other continents (Figure 2B).

**Figure 2.**
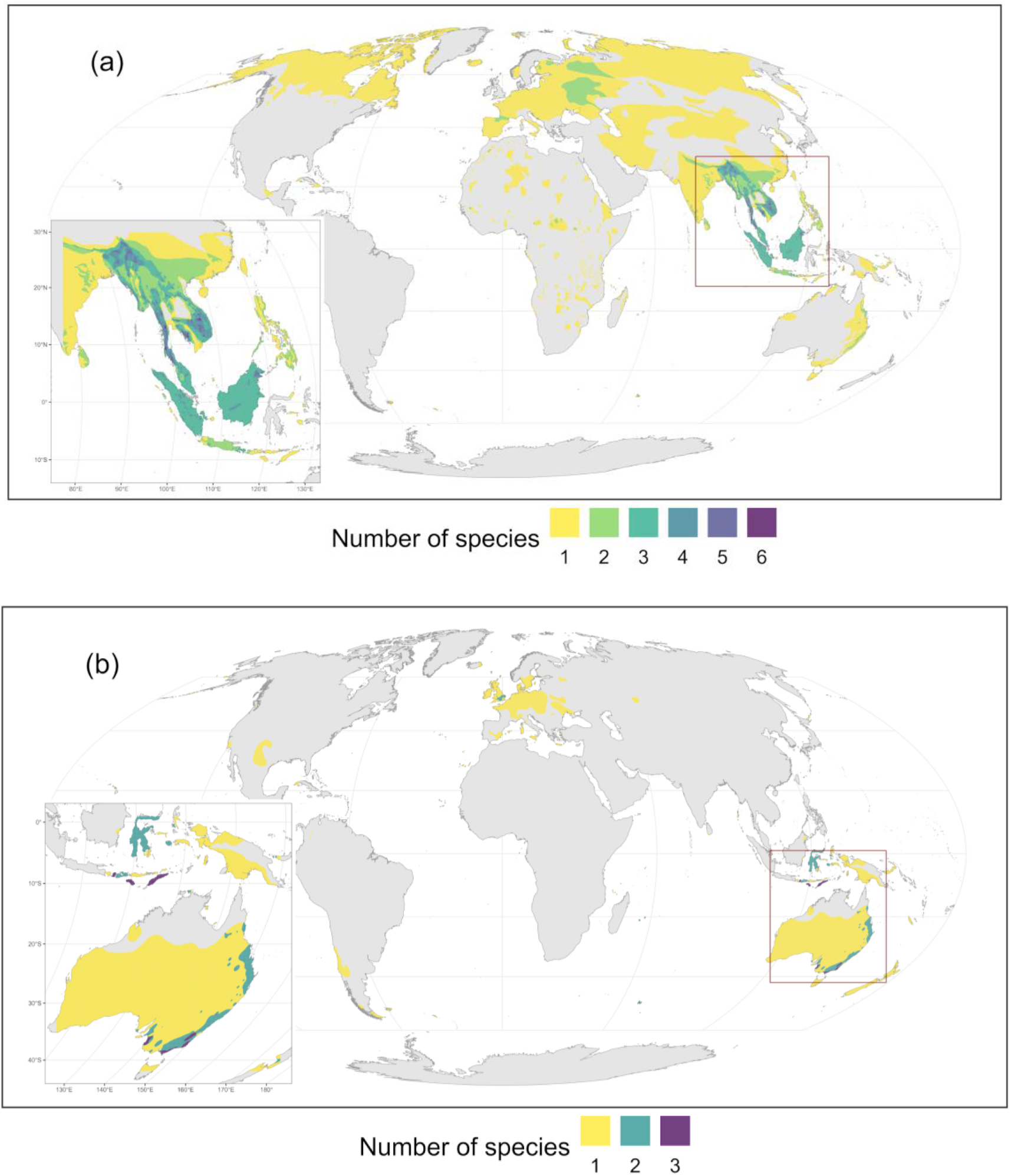
Distribution of (a) the native and (b) alien ranges of alien threatened mammals (n = 41).

When investigating the continental flows of alien threatened mammals from their native to their alien ranges, we found that the intra-continental flows (i.e., species establishment in other parts of the same continent) were particularly pronounced (Figure 3). Most intra-continental exchanges occurred within Asia (n = 15), Oceania (n = 7), and Europe (n = 5; Figure 3). Contrarily, the major inter-continental flows happened from Asia to Oceania (n = 7), from Asia to Europe (n = 5), and from Asia to North America (n = 4; Figure 3).

**Figure 3.**
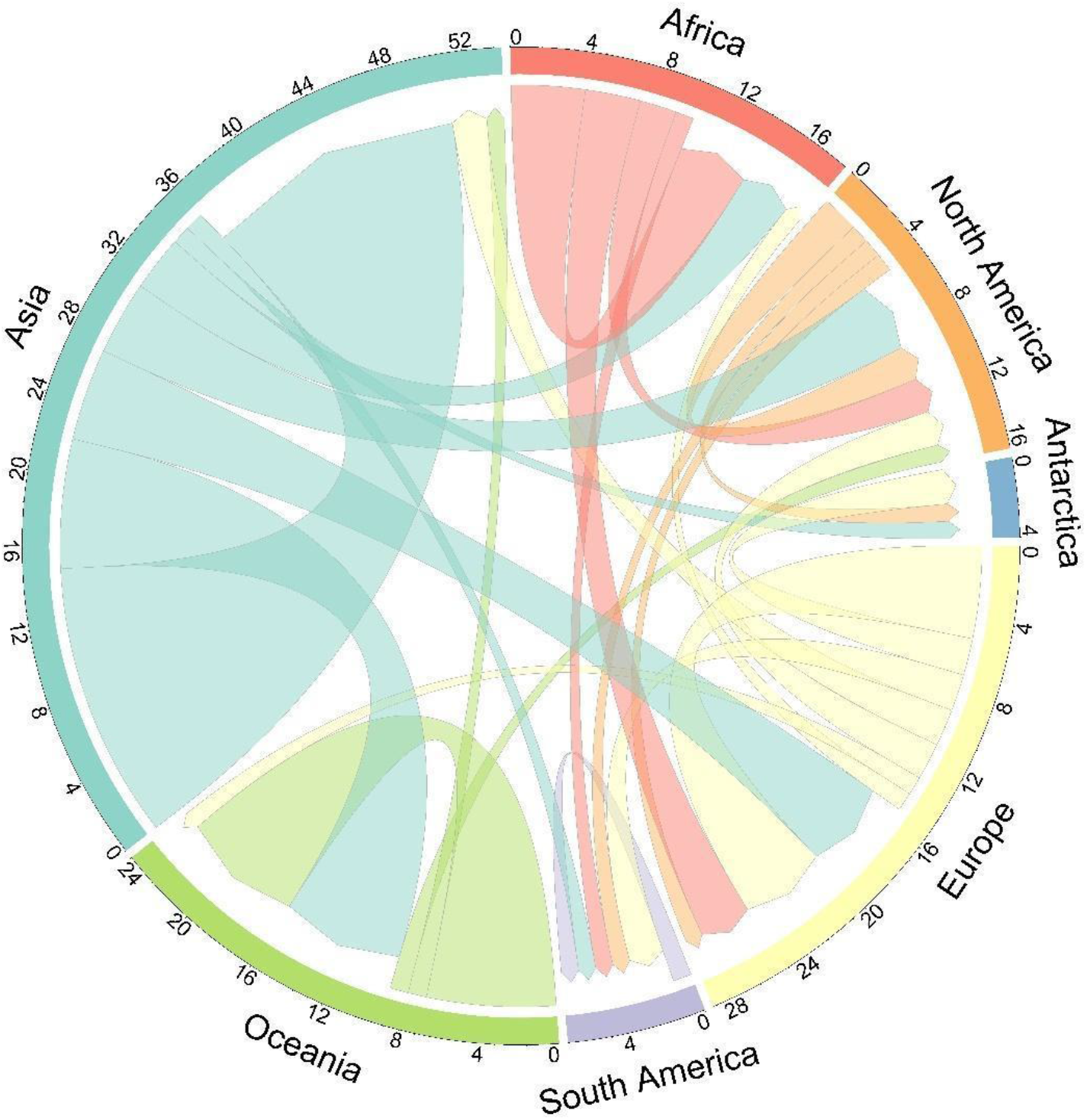
Continental flows of the 41 alien threatened mammals between donor and receiver continents. The width of the lines indicates the number of species exchanged between donor and receiver continents. Multiple species’ introductions to or originating from the same continent are counted once. Links attached closer to the circle base indicate species native from that continent and introduced to the continent where the link ends further and with an arrow.

The most important introduction pathways of alien threatened mammals were hunting (n = 94), followed by farming (n = 38), and pet trade (n = 27; Supplementary Results).

### Causes of threats and conservation measures

All alien threatened mammals are affected by more than one threat. The dominant threat is biological resource use (n = 67), followed by agriculture and aquaculture (n = 63), and invasive species, genes and diseases (n = 46; Figure 4A). Native populations of the study species are subjected to a range of different conservation measures, the most important being species management (n = 52), followed by land/water management and land/water protection (both n = 43; Figure 4B).

**Figure 4.**
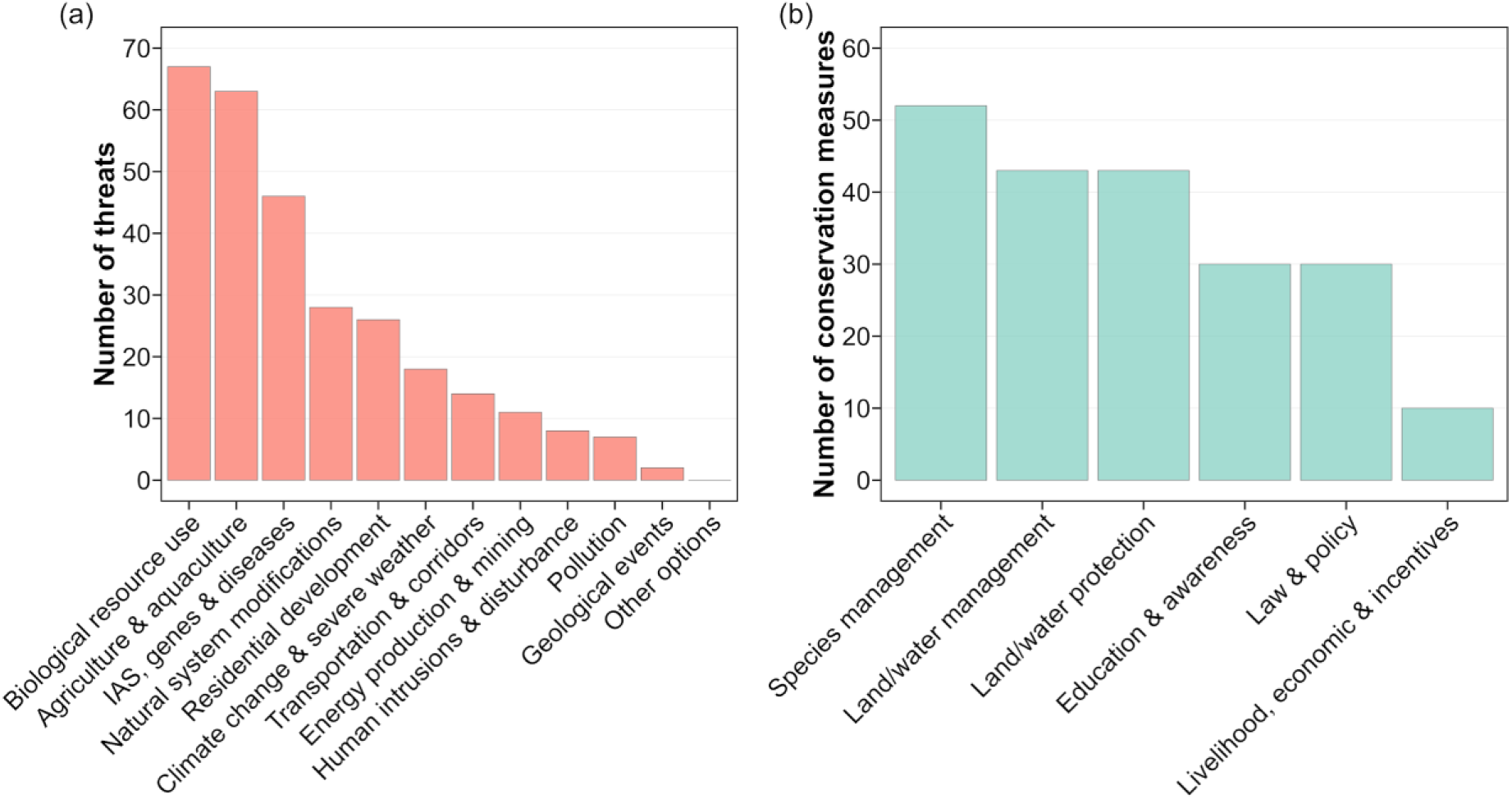
The Level 1 (a) threats to alien threatened mammals and (b) conservation measures applied to populations in the native ranges of alien threatened mammals (n = 41 species). Each species can be affected by more than one threat or be subjected to more than one conservation measure. The shortened Level 1 threats are indicated as follows: Residential & commercial development: “Residential development”, Transportation & service corridors: “Transportation & corridors”, Invasive & other problematic species, genes & diseases: “IAS, genes & diseases”. The shortened Level 1 conservation measure is indicated as follows: Livelihood, economic & other incentives: “Livelihood, economic & incentives”.

### Including alien populations in global extinction risk assessments

We found that including alien populations into the study species’ global extinction risk assessment resulted in changes of extinction risk categories of 9 (22%) of the 41 alien threatened mammals studied. Three species changed from CR to EN, one species from EN to VU, one species from EN to Least Concern (LC), two species from VU to Near Threatened (NT), and two species from VU to LC (Supplementary Results, Figure 6). The most notable changes occurred when a study species shifted its classification by two or more levels. Those include the European rabbit (*Oryctolagus cuniculus*), which shifted from EN to LC, and the Javan (*Rusa timorensis*) and Sambar deer (*R. unicolor*), both transitioning from VU to LC. All these species have been introduced in several continents, and many of their alien populations have become invasive. Often those alien populations are showing high population numbers and positive trends, thereby notably increasing the number of mature individuals, or lessening the population reduction in the native range. Lastly, the calculated RLI for the original assessments was 0.47, while the RLI for our re-assessments was 0.53.

**Figure 6.**
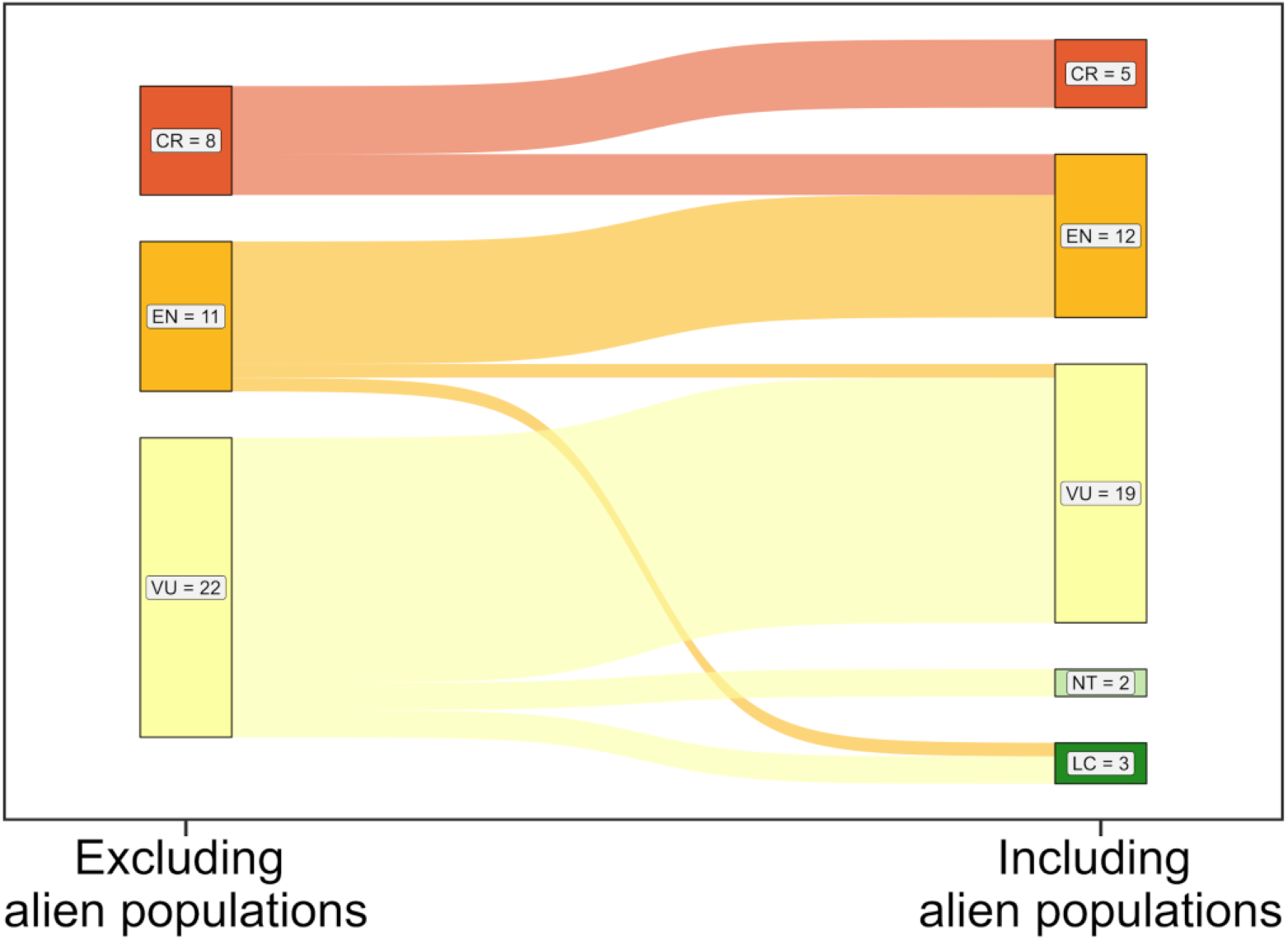
Changes in global IUCN Red List assessments based on original assessments (on the left; IUCN 2023) and our assessment (including alien populations, on the right) of alien threatened mammals (n = 41 species). Red List Categories are indicated with their code: CR (Critically Endangered), EN (Endangered), VU (Vulnerable), Near Threatened (NT), and Least Concern (LC). Assessments are grouped by IUCN Red List Category (CR, EN, VU, NT, LC). Flow colors refer to Red List Categories (red for CR, orange for EN, yellow for VU, light green for NT, forest green for LC). For details, see Supplementary Results.

## DISCUSSION

### Diversity and distributions of alien threatened mammals

In our study, we found 41 alien threatened mammals, a higher number than what previously reported in the study of Gibson & Yong (2017) (19 species). Their analysis excluded all threatened mammals introduced as benign introductions, whereas our analysis considered species introduced through this pathway for which alien populations were not covered in Red List assessments (or with an unclear inclusion status). Moreover, a comprehensive database on alien mammals’ distributions was recently released (Biancolini et al. 2021). In the 20 years that passed from the publication of Long (2003), from which Gibson & Yong (2017) take their information, many more species have been recognized as aliens–including severalthreatened mammals with alien populations (e.g., the Mexican black agouti *Dasyprocta mexicana*). Furthermore, some mammals listed by Gibson & Yong (2017) are either not threatened anymore (e.g., the greater stick-nest rat *Leporillus conditor*) or do not have alien populations in DAMA (e.g., the Arabian oryx *Oryx leucoryx*; Biancolini et al. 2021)–for instance, because the alien population was eradicated or is not viable anymore, or because of a re-evaluation of the populations’ status.

Many of our study species are native to areas where it is difficult to assess their threat level and the causes of population declines. Indeed, 14% of all mammals are categorized as Data Deficient in the Red List (IUCN 2023) highlighting that more research is needed to carefully evaluate threat levels in the native range, which is essential to make informed decisions especially considering rapid global changes and increased extinction and introduction rates.

### The importance of intra-continental introductions

Intriguingly, a significant number of threatened mammals have been introduced near the native range on the same continent, as happened several times in Asia and Oceania. For conservation purposes, similar to benign introductions, populations of the species are introduced and established—often in proximity to their native range—in a different location to create a wild population. This happens often in areas where the species is less likely to face the same threats, or where threat intensity is lower. Importantly, Red List guidelines (IUCN 2022a) refer to introductions within the taxon’s current natural range, but they do not refer to historic ranges. In principle, introductions within the historic range may not be considered alien populations, and this could reduce the pool of study species (but also the global pool of alien mammals). Unfortunately, historic ranges are difficult to obtain, and to date there is no such information for mammals.

Despite several conservation introductions, similar to Gibson & Yong (2017), we found that populations of threatened mammals were primarily introduced for hunting purposes. Notably, the top threat (biological resource use, such as bushmeat or trophy hunting) pushing our study species towards extinction is also the same major pathway that created their alien ranges (hunting). The impact of hunting on mammal populations has been staggering, resulting in a population reduction of more than 80% since 1970 (Benítez-López et al. 2017). Hunting was the prevalent pathway of historical introductions, whereas recent introductions are more driven by pet trade (Biancolini et al. 2021, Tedeschi et al. 2022). Notably, recently mammals’ introductions have declined sharply (Seebens et al. 2017). The most common conservation approach for threatened mammals involves their management, with many species specifically protected through harvest management, underlying the effort in place to protect them from overexploitation.

### Beyond categories: how the inclusion of alien populations reshapes Red List assessments

Including the introduced populations in Red List assessments can have several implications. On the one hand, if this inclusion decreases the Red List category, protection and conservation efforts towards species’ native populations may decrease, as well as public awareness and funding. Conversely, thriving but monitored alien populations may, under specific circumstances (see below), be considered for conservation actions.

We excluded from the analysis 12 species of threatened mammals whose populations were already included in Red List assessments, but interesting is the case of introductions driven by conservation goals but not included in the assessments. This seems to be the case for the European mink (*Mustela lutreola*), for the Northern quoll (*Dasyurus hallucatus*) and for the Tasmanian devil (*Sarcophilus harrisii*), while it is unclear if it happened for the woylie (*Bettongia enicillate*), the Aders’ duiker (*Cephalophus adersi*), the Balabac mouse deer (*Tragulus nigricans*) and for the koala (*Phascolarctos cinereus*) as well.

In a couple of cases this is the result of outdated Red List assessments. For instance, the European mink was last assessed in 2015 and Maran et al. (2016) report no established populations on the Kuril Islands in 2014. However, evidence of presence is reported on the islands for the period 2014-2021 (Biancolini et al. 2021, Kisleyko et al. 2021). Introductions were sometimes mentioned in Red List assessments but were then omitted from the geographic range map (as in the case of the woylie; Woinarski & Burbidge 2016) or from the “conservation actions” section (as for the Aders’ duiker; IUCN SSC Antelope Specialist Group 2017), challenging the understanding of the alien populations’ inclusion in the assessments.

Moreover, in some cases the unavailability of information hindered the possibility of re-assessing the study species. The Mexican black agouti was classified CR in 2008 (Vázquez et al. 2008), and it has recently been reported to be invasive in Cuba (Borroto-Páez & Mancina 2017). This possibly implies that the population is either stable or increasing, but without knowing the introduced population’s trend, we can only speculate on its conservation relevance. In those cases, an updated assessment of the species is essential to make informed management and conservation decisions.

For other study species, the introduced population is present in an area that is too small to sustain an adequate number of individuals (e.g., the dusky pademelon *Thylogale brunii* introduced on the 400 km^2^ of Kai Kecil Island; Leary et al. 2016), or it is known that the abundance of the alien population is low, as for the 21 alien individuals of Balabac mouse deer (Widmann 2015). In those cases, the conservation relevance of the introduced population is likely low.

In some instances, the introduced population already made (or could make) a difference, as underlined by the moderate increase in the RLI calculated on our re-assessments. The Celebes’ crested macaque’s (*Macaca nigra*) introduced population in Indonesia probably exceeds that in its native range, and it is likely facing less threats than in the native range (where it is a favored bushmeat species; Hilser et al. 2013, Lee et al. 2020). The Australian introduced population of banteng (*Bos javanicus*) is reported to be thriving and likely not facing the same threats as in the native range (Gardner et al. 2016). Lastly, some study species have already been in the spotlight because of the conservation paradox they represent, such as the aoudad (*Ammotragus lervia*; Cassinello 2018) or the European rabbit (Lees & Bell 2008). For all those species, some introduced populations could act as a backup, an “ark” for future conservation actions (Gibson & Yong, 2017) or as “safety populations”, which could avoid the extinction in the wild in case of abrupt and drastic native population declines. Importantly, alien populations of threatened species should not be exempted from monitoring or management programs. Current and future impacts should be strictly monitored, as they can arise after a time-lag, a phenomenon called “invasion debt” (Essl et al. 2011). If the invasion stage is reached, control and eradication are mandatory, possibly with translocation of individuals to captivity, which avoids specimens’ depletion (a common *ex-situ* conservation drawback; Snyder et al. 1996, Conde et al. 2011) while at the same time preventing them to adversely impact biotas. Above all, a thorough impact assessment is crucial to determine whether alien populations can contribute to species conservation and can, for instance, be included in a safe list—guiding conservationists on which taxa can be used safely, if necessary, without promoting biological invasions (Kumschick et al. 2024).

Captive breeding and reintroductions using alien populations have already been suggested as potential conservation measures (Gibson & Yong, 2017). While caution is advised, as alien populations can harbor parasites or be genetically impoverished (Gibson & Yong, 2017), using them *in-situ* can provide benefits such as increased genetic diversity, avoidance of adaptations to captivity, and lower costs (Conde et al. 2011) while excluding or minimizing the threats in the native range. Additionally, conserving species close to the native range region, as in the majority of alien threatened mammals, is usually advised (Conde et al. 2011, Pritchard et al. 2012).

## ACKNOWLEDGEMENTS

FE and AS appreciate funding by the Austrian Science Foundation FWF [Project DOIs: 10.55776/I5825; 10.55776/P34688].

