## Supplementary Information for "A synthesis on alien mammals threatened in their native range"

**Supplementary Methods**

Taxonomy

In our study we followed the taxonomy provided by IUCN (IUCN 2023). When comparing the list of native threatened mammals (IUCN 2023) to the list of alien mammals (Biancolini et al. 2021) we found the following inconsistencies. Nine species (of genera *Macropus, Microtus*, and *Myodes*, plus the mouflon, *Ovis orientalis*) were reclassified recently, and the taxonomy of DAMA is different from the taxonomy of IUCN. However, all of those species are LC or NT, and therefore are in any case excluded from our study. Three species (water buffalo, camel, and llama) are present in DAMA, but IUCN does not assess them (nor map their distribution) as they are considered domestic species and are therefore excluded from our study. Lastly, one species (arctic fox) does not have distribution data in IUCN, but it's assessed as LC and is excluded from our study.

Table S1: Summary of the criteria used by IUCN to classify species as Vulnerable (VU), Endangered (EN), Critically Endangered (CR), or Extinct in the wild (EW).

| **Category** | **Criteria** |
|---|---|
| Vulnerable (VU) | A species is considered VU, EN, or CR when the best available evidence indicates that it meets any of the criteria A to E for, respectively, the categories VU, EN, or CR, and it is therefore considered to be facing a high risk of extinction in the wild. |
| Endangered (EN) | |
| Critically endangered (CR) | |
| Extinct in the wild (EW) | A species is EW when it is known only to survive in captivity or as an alien, and it is presumed to be EW when exhaustive surveys in known and/or expected habitat, at appropriate times, throughout its historic range have failed to record any specimens. |

Table S2: Taxonomic species' names used in IUCN, in DAMA, common names, and reason to not include the species in our study.

| **IUCN Taxonomy** | **DAMA Taxonomy** | **Common name** | **Rationale for exclusion from the study** |
|---|---|---|---|
| NA | *Bubalus bubalis* | Water buffalo | *Bubalus arnee* is the wild water buffalo, while *Bubalus bubalis* is not present in IUCN |

| | | | |
|---|---|---|---|
| | | | because it's the domestic form. |
| NA | *Camelus dromedarius* | Camel | Domesticated, thus not present in IUCN. |
| NA | *Lama glama* | Llama | Domesticated, thus not present in IUCN. |
| *Notamacropus agilis* | *Macropus agilis* | Agile wallaby | Classified as LC in IUCN. |
| *Notamacropus eugenii* | *Macropus eugenii* | Tammar wallaby | Classified as LC in IUCN. |
| *Notamacropus parma* | *Macropus parma* | Parma wallaby | Classified as NT in IUCN. |
| *Notamacropus rufogriseus* | *Macropus rufogriseus* | Red-necked wallaby | Classified as LC in IUCN. |
| *Microtus mystacinus* | *Microtus levis* | East European vole | Classified as LC in IUCN. |
| *Clethrionomys gapperi* | *Myodes gapperi* | Southern Red-backed vole | Classified as LC in IUCN. |
| *Clethrionomys glareolus* | *Myodes glareolus* | Bank vole | Classified as LC in IUCN. |
| *Clethrionomys rutilus* | *Myodes rutilus* | Northern Red-backed vole | Classified as LC in IUCN. |
| *Ovig gmelini* | *Ovis orientalis* | Mouflon | Classified as NT in IUCN. |
| *Vulpes lagopus* | *Vulpes lagopus* | Arctic fox | Distribution range not mapped, but species classified as LC in IUCN. |

Alien threatened mammals range visualization

For visualization purposes, we removed native species range polygons where alien threatened mammals are extinct or with uncertain presence, and those where they are introduced, vagrant, have an uncertain origin, or originate from benign introductions (i.e., species' established populations, for conservation purposes, outside its current native range and within a suitable eco-geographical area and habitat; IUCN/SSC 2013, IUCN 2022a), keeping the DAMA ranges.

Hierarchical structure of threats and conservation measures

IUCN provides a hierarchical structure of causes of threats and applied conservation measures composed by three levels, and assessors are asked to indicate the threats/conservation measures that triggered the listing of the species at the lowest level possible (IUCN 2016). The adoption of a level can be interpreted as a measure of accuracy in the assessment Assessors make of the threats/conservation measures to the species. The more certain information they have, the more likely they chose a lower (i.e., more specific) hierarchical level. Here, we aggregate all threats and conservation measures at lower levels to the corresponding highest level (Level 1).

IUCN provides a hierarchical structure of threat types, including 12 major threats: 1) Residential & commercial development, 2) Agriculture & aquaculture, 3) Energy production & mining, 4) Transportation & service corridors, 5) Biological resource use, 6) Human intrusions & disturbance, 7) Natural system modifications, 8) Invasive & other problematic species, genes & diseases, 9) Pollution, 10) Geological event, 11) Climate change & severe weather, and 12) Other options (IUCN 2022a). Those major threats are further divided in sub-categories, for instance 2) Agriculture & Aquaculture is divided into 2.1.) Annual & perennial non-timber crops, 2.2.) Wood & pulp plantations, etc. (IUCN 2022a). These are divided again into other sub-categories, for example 2.1.) Annual & perennial non-timber crops are further divided into 2.1.1.) Shifting agriculture, 2.1.2.) Small-holder farming, and so on (IUCN 2022a). For simplicity, we refer to "Level 1 threats", "Level 2 threats", and "Level 3 threats" throughout.

Among the Level 1 threat category 8) Invasive & other problematic species, genes & diseases, we retained the following Level 2 and Level 3 threats: invasive non-native/alien species/diseases, introduced genetic material, problematic species/diseases of unknown origin, and diseases of unknown cause. We assumed a species/disease of unknown origin or cause to be introduced.

Continental flows visualization

For this visualization, each study species native or introduced to a continent was considered only once, regardless of the number of times the species has been considered native in or introduced to that continent.

IUCN Red List re-assessment

The IUCN Red List categorization process should also be applied to wild subpopulations resulting from introductions outside the natural range (also termed benign introductions or translocations), if all of the following conditions are met (IUCN/SSC 2013, IUCN 2022a):

(a) The known or likely intent of the introduction was to reduce the extinction risk of the taxon being introduced.

(b) The introduced subpopulation is geographically close to the natural range of the taxon. What is considered to be geographically close enough is determined by the assessor.

(c) The introduced subpopulation has produced viable offspring (i.e., offspring that have reached maturity or are likely to do so).

(d) At least five years have passed since the introduction.

Moreover, those criteria should be applied as well to established (i.e., self-sustaining) translocated or re-introduced subpopulations (within the taxon's natural range), regardless of the original goal of such translocations or re-introductions (IUCN/SSC 2013, IUCN 2022a).

Red List Index (RLI)

The RLI shows trends in the status of taxa (Butchart et al. 2007): a RLI value of 1 signifies that all species are categorized as LC, indicating they are not expected to become EX in the foreseeable future. Conversely, a RLI value of 0 indicates the scenario where all species have become EX. If the RLI value remains constant over time, it suggests that the overall extinction risk for the group has not changed.

Full list of R packages used for statistical analysis and data visualization

To test if the proportion of alien threatened mammals statistically differed from the proportion of alien or threatened mammals while accounting for non-independence in the observations, we used McNemar's exact test (McNemar 1947) from the R package *exact2x2* v. 1.6.9 (Fay 2010). We used *rredlist* v. 0.7.1 (Gearty & Chamberlain 2022) to extract species' Red List category, countries of occurrence, threats, and conservation measures. We assigned the continents of occurrence to each species by intersecting its native and alien countries of occurrence with *rnaturalearth* v. 0.1.0 (South 2017) continents. We used *circlize* v. 0.4.15 (Gu 2014) to produce chord diagrams of species' flows between continents (see also "Continental flows visualization").

**Supplementary Results**

Table S3: The 53 alien mammals threatened in their native ranges we first identified. Order, family, scientific name, and IUCN threat category are given. Species not included in the study (because they are benign introductions and thus are already included in Red List assessments) are marked with an asterisk (*) and are written in grey font.

| **Order** | **Family** | **Scientific name** | **Common name** | **Category** |
|---|---|---|---|---|
| Carnivora | Mustelidae | *Mustela lutreola* | European Mink | CR |
| Artiodactyla | Bovidae | *Ammotragus lervia* | Aoudad | VU |
| | | *Beatragus hunteri** | Hirola | CR |
| | | *Bos javanicus* | Banteng | EN |
| | | *Cephalophus adersi* | Aders' Duiker | VU |
| | | *Gazella subgutturosa* | Goitered Gazelle | VU |
| | | *Nanger soemmerringii* | Soemmerring's Gazelle | VU |
| | | *Tragelaphus derbianus* | Giant Eland | VU |
| | Cervidae | *Axis porcinus* | Hog Deer | EN |
| | | *Hydropotes inermis* | Chinese Water Deer | VU |
| | | *Rangifer tarandus* | Reindeer | VU |
| | | *Rusa marianna* | Philippine Deer | VU |
| | | *Rusa timorensis* | Javan Deer | VU |
| | | *Rusa unicolor* | Sambar Deer | VU |
| | Hippopotamidae | *Hippopotamus amphibius* | Hippopotamus | VU |
| | Suidae | *Babyrousa babyrussa* | Babirusa | VU |
| | Tragulidae | *Tragulus nigricans* | Balabac Mouse Deer | EN |

| | | | | |
|---|---|---|---|---|
| Dasyuromorphia | Dasyuridae | *Dasyurus hallucatus* | Northern Quoll | EN |
| | | *Parantechinus apicalis** | Dibbler | EN |
| | | *Sarcophilus harrisii* | Tasmanian Devil | EN |
| Diprotodontia | Macropodidae | *Dendrolagus matschiei* | Huon Tree-Kangaroo | EN |
| | | *Lagorchestes hirsutus** | Rufous Hare Wallaby | VU |
| | | *Lagostrophus fasciatus** | Banded Hare Wallaby | VU |
| | | *Petrogale lateralis** | Pearson Island Rock-Wallaby | VU |
| | | *Petrogale penicillata* | Brush-Tailed Rock Wallaby | VU |
| | | *Petrogale persephone** | Proserpine Rock Wallaby | EN |
| | | *Thylogale browni* | New Guinea Pademelon | VU |
| | | *Thylogale brunii* | Dusky Pademelon | VU |
| | Phascolarctidae | *Phascolarctos cinereus* | Koala | VU |
| | Potoroidae | *Bettongia penicillata* | Woylie | CR |
| | | *Potorous gilbertii** | Gilbert's Potoroo | CR |
| Eulipotyphla | Talpidae | *Desmana moschata* | Russian Desman | CR |
| Lagomorpha | Leporidae | *Lepus corsicanus* | Corsican Hare | VU |
| | | *Oryctolagus cuniculus* | European Rabbit | EN |
| Pholidota | Manidae | *Manis culionensis* | Philippine Pangolin | CR |
| Primates | Callitrichidae | *Saguinus oedipus* | Cotton-Top Tamarins | CR |
| | Cercopithecidae | *Macaca arctoides* | Stump-Tailed Macaque | VU |

| | | | | |
|---|---|---|---|---|
| | | *Macaca fascicularis* | Nicobar Crab-eating Macaque | EN |
| | | *Macaca leonina* | Northern Pig-Tailed Macaque | VU |
| | | *Macaca nemestrina* | Southern Pig-tailed Macaque | EN |
| | | *Macaca nigra* | Celebes Crested Macaque | CR |
| | | *Macaca sylvanus* | Barbary Ape | EN |
| | | *Piliocolobus kirkii** | Zanzibar Red Colobus | EN |
| | | *Trachypithecus auratus* | Javan Lutung | VU |
| | Daubentoniidae | *Daubentonia madagascariensis** | Aye-aye | EN |
| | Lemuridae | *Eulemur albifrons* | White-Fronted Lemur | VU |
| | | *Eulemur fulvus* | Brown Lemur | VU |
| | | *Eulemur mongoz* | Mongoose Lemur | CR |
| | | *Varecia variegata** | Black-and-white Ruffed Lemur | CR |
| Proboscidea | Elephantidae | *Elephas maximus* | Asian Elephant | EN |
| Rodentia | Capromyidae | *Geocapromys ingrahami** | Bahaman Hutia | VU |
| | Dasyproctidae | *Dasyprocta mexicana* | Mexican Black Agouti | CR |
| | Muridae | *Pseudomys fieldi** | Djoongari | VU |

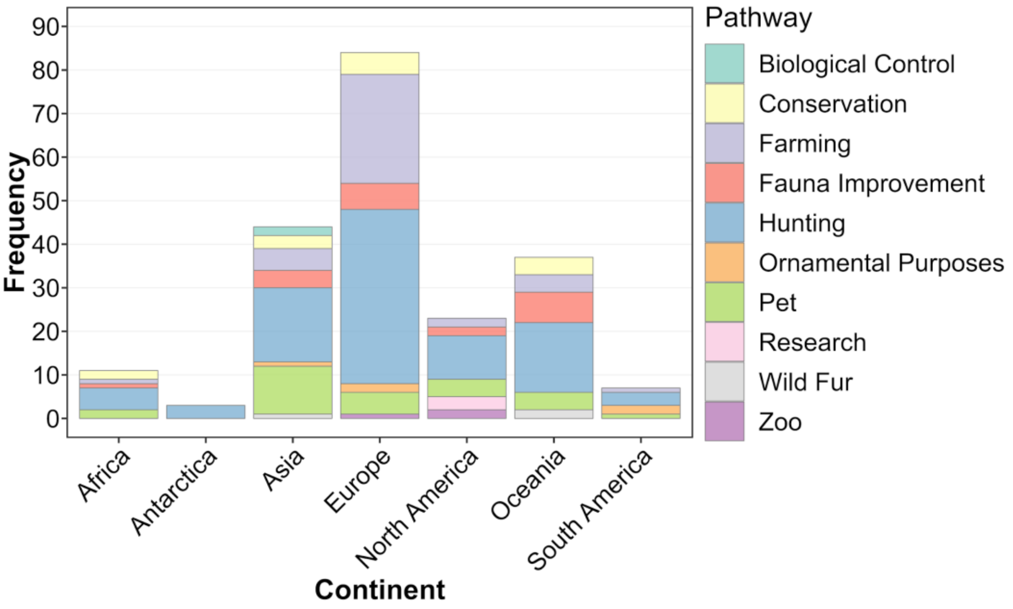


Figure S1: Pathways of introduction of alien threatened mammals (n = 41 species). The same species could have been introduced to the same continent through different pathways, and vice versa (i.e., the same species could have been introduced through the same pathway into different continents). The pathway "conservation" refers to introductions performed for conservation purposes not based on IUCN Guidelines (IUCN/SSC 2013, IUCN 2022b).

<u>Considering alien populations in global extinction risk assessments</u>

*CRITICALLY ENDANGERED*

*Mustela lutreola*

There is an introduced population for conservation purposes on Kuril Islands (Russian Federation), which is not included in Red List assessment (Maran et al. 2016). A pers.comm of 2014 available in the assessment tells that population is not established. Kisleyko et al. (2021) found 36 occurrences of *M. lutreola* on Kuril Islands from 2014 to 2021. As, in comparison, *M. lutreola* has high numbers in the native range, even adding those few individuals present on Kuril Islands the assessment likely would not change. We do not estimate a change in IUCN Red List Category.

*Bettongia penicillata*

Introduced as a benign introduction, but it is unclear from Red List assessments if the species is included (Woinarski & Burbidge 2016a). Low reintroduction success rate, but no useful assessment information on the status of the three populations on the South Eastern islands (Australia) indicated in DAMA (Biancolini et al. 2021). Probably those populations are declining as well (as the native ones on the mainland). We do not estimate a change in IUCN Red List Category.

*Desmana moschata*

There is no information available (Rutovskaya et al. 2023). We do not estimate a change in IUCN Red List Category.

*Manis culionensis*

Contrasting evidence for presence of the species on Balabac Island (Philippines; Schoppe et al. 2019). Introduced and present on Apuli Island (Philippines), but no information on population trend is available (Schoppe et al. 2019; Biancolini et al. 2021). In any case, the surface of Apuli Island is likely too small to sustain an adequate number of individuals. We do not estimate a change in IUCN Red List Category.

*Eulemur mongoz*

Given that the population on the Comoros Islands is probably larger (Razafindramanana et al. 2020) and less at risk than the one in Madagascar, including the alien population would lower the classification to at least EN if not VU. One would need to know the population trend on the Comoros Islands. We do not estimate a change in IUCN Red List Category.

*Macaca nigra*

The alien population on Bacan Island (2800 $km^2$; Indonesia) is not included in the IUCN RL assessment (Lee et al. 2020). It's not a benign introduction (Biancolini et al. 2021), and the population likely undergoes fewer human pressures (e.g., hunting) compared to the native range, and it has high numbers (Hilser et al. 2013). There were 100,000 individuals in the late 1990s. Considering the number of introduced individuals, the IUCN categorization of the species would certainly diminish. Adding also Bacan Island's area to *M. nigra* Extent Of Occurrence (EOO), it would pass the threshold of 100$km^2$ (required to be categorized as CR under criterion B1). With information on the population trend in the native range, the categorization could shift even more. We estimate a change in IUCN Red List Category to EN.

*Saguinus oedipus*

The introduced population in Colombia is still present, but it is not considered in the IUCN RL assessment (Rodríguez et al. 2021). Probably nothing would change as the locations are very small and there is no information on the numbers of individuals. We do not estimate a change in IUCN Red List Category.

*Dasyprocta mexicana*

Assessed as CR in 2008 (Vázquez et al. 2008), reported as being invasive in Western Cuba (Borroto-Páez & Mancina 2017). If the species is classified as invasive, it must have enough individuals to produce negative impacts, and the population must still be growing or at least be stable. Thus, the classification could shift to EN, VU, or even NT. We estimate a change in IUCN Red List Category to EN.

*ENDANGERED*

*Axis porcinus*

The introduced populations (several, scattered across Australia, Andamane and Nicobare Islands, possibly Sri Lanka; Biancolini et al. 2021) are not included in the assessment (Timmins et al. 2015a), but as they are declining, it is likely that the inclusion wouldn't change the assessment. We do not estimate a change in IUCN Red List Category.

*Bos javanicus*

The IUCN RL assessment indicates 4000-8000 wild native reproductive individuals, but Garik Gunak Barlu National Park (Australia) population (one of the several introduced populations of the species) alone has 6000 total individuals (Gardner et al. 2016). The Australian population is thriving and genetically it's composed by wild individuals (Gardner et al. 2016), and it's highly likely that it's not experiencing the same pressures as of the native populations. Therefore, if the Australian population would be considered in the assessment, the species would likely have a lower threat category. We estimate a change in IUCN Red List Category to VU.

*Tragulus nigricans*

Only 21 individuals have been reported to be present on Calauit Island (Philippines; Widmann 2015). Likely irrelevant. We do not estimate a change in IUCN Red List Category.

*Dasyurus hallucatus*

Introduced as benign introduction (Biancolini et al. 2021) but not included in Red List assessment (Oakwood et al. 2016). Introduced on two Australian islands so small that it is likely irrelevant. We do not estimate a change in IUCN Red List Category.

*Sarcophilus harrisii*

Introduced as a benign introduction (Biancolini et al. 2021), but the assessment is very old (2008; Hawkins et al. 2008) and it actually seems to not include the introduced populations. It was introduced on Maria Island (Australia), but the island is too small, and the introduced population is likely in semi-captivity. We do not estimate a change in IUCN Red List Category.

*Dendrolagus matschiei*

Population estimates of 2500 mature individuals, EOO 14000 $km^2$ (Ziembicki & Porolak 2016). Adding the introduced populations of Umboi Island (930 $km^2$; Papua New Guinea) and the Western part of New Britain Island (more than 1000 $km^2$; Papua New Guinea) wouldn't allow to pass the 20000 $km^2$ threshold required to shift from EN to VU. The number of mature individuals would be higher than 2500 (which could cause a shift to VU). But precise population trends are difficult to obtain for this species. We do not estimate a change in IUCN Red List Category.

*Oryctolagus cuniculus*

There are so many introduced populations scattered globally (Biancolini et al. 2021) that including them would clearly shift the categorization to LC. Several of those populations are invasive and causing environmental and socio-economic negative impacts (Villafuerte & Delibes-Mateos 2019). Some introduced populations that are not invasive could be safeguarded as "safety populations". We estimate a change in IUCN Red List Category to LC.

*Macaca fascicularis*

Even after removing the pre-historical introductions (several across Indonesia), the areas where the introduced populations are present are small and scattered (Biancolini et al. 2021, Hansen et al. 2022). Likely irrelevant. We do not estimate a change in IUCN Red List Category.

*Macaca nemestrina*

Some of the introduced populations in DAMA (e.g., Singapore; Biancolini et al. 2021) are not established/viable (Ruppert et al. 2022). Very small areas scattered across the globe (from Indonesia to Cuba) remain (Biancolini et al. 2021). Likely irrelevant. We do not estimate a change in IUCN Red List Category.

*Macaca sylvanus*

The alien population in Gibraltar is very small (300 individuals) so it is likely irrelevant (Wallis et al. 2020). But the alien population is stable and could be valuable in the future. We do not estimate a change in IUCN Red List Category.

*Elephas maximus*

The species is considered native (genetic evidence; Williams et al. 2020) in the alien range indicated in DAMA (Borneo Island; Malesia, Indonesia, Brunei; Biancolini et al. 2021), therefore the available information does not allow a complete re-assessment. We do not estimate a change in IUCN Red List Category.

*VULNERABLE*

*Ammotragus lervia*

As it is assessed with criterion C1 (Cassinello et al. 2022), considering also the alien populations (especially the Spanish one and North American one) probably could shift the category to NT or even LC, as the mature individuals would be more than 10,000 and likely not decreasing (as some of those are invasive populations that could be stable or increasing). We estimate a change in IUCN Red List Category to NT.

*Babyrousa babirussa*

Prehistoric introduction on Buru Island (Indonesia) already included in the IUCN RL assessment (Macdonald et al. 2008). We do not estimate a change in IUCN Red List Category.

*Cephalophus adersi*

Introduced only on two very small islands in Madagascar (5 individuals introduced in one of them; IUCN SSC Antelope Specialist Group 2017, Biancolini et al. 2021). Likely irrelevant. We do not estimate a change in IUCN Red List Category.

*Gazella subgutturosa*

Introduced for conservational purposes (Biancolini et al. 2021), it seems the alien population is already included in the assessment (IUCN SSC Antelope Specialist Group 2017b). Wide native distribution, introduced on two very small islands (in the Middle East) too small to make a difference. Likely irrelevant. We do not estimate a change in IUCN Red List Category.

*Hippopotamus amphibius*

Too few individuals in the introduced range (Colombia; Lewison & Pluháček 2017, Biancolini et al. 2021). Likely irrelevant. We do not estimate a change in IUCN Red List Category.

*Hydropotes inermis*

The introduced population (United Kingdom; Biancolini et al. 2021) occupies an extended area, and the population numbers are relatively good (Harris & Duckworth 2015). It could potentially be a good addition, and especially it likely doesn't have the same pressures (poaching and habitat destruction) that the species has in the native range. We estimate a change in IUCN Red List Category to NT.

*Nanger soemmerringii*

The introduced population on Dahlak Kebir and Norah Islands (Eritrea) is the bigger one (IUCN SSC Antelope Specialist Group 2016). Without considering it, the species would be EN or CR. From the IUCN RL assessment it is not clear if the introduction has conservational purposes (in that case, it could be included in the assessment; IUCN SSC Antelope Specialist Group 2016). We do not estimate a change in IUCN Red List Category.

*Rangifer tarandus*

Wide native distribution with many individuals (Gunn 2016), introduced populations (small islands scattered across the globe; Biancolini et al. 2021) too small to make a difference. Likely irrelevant. We do not estimate a change in IUCN Red List Category.

*Rusa marianna*

There is no information available (MacKinnon et al. 2015). We do not estimate a change in IUCN Red List Category.

*Rusa timorensis*

Assessed as VU because there are less than 10,000 wild native mature individuals (Hedges et al. 2015). The introduced populations in several areas of the world have high numbers (120,000 in New

Caledonia, 60,000 in Mauritius, etc; Hedges et al. 2015). Several populations are invasive and are causing negative impacts (Hedges et al. 2015). Some introduced populations that are not invasive could be safeguarded as “safety populations”. We estimate a change in IUCN Red List Category to LC.

*Rusa unicolor*

Introduced populations in several areas of the world (Biancolini et al. 2021). Several populations are invasive and are causing negative impacts (Timmins et al. 2015b). Some introduced populations that are not invasive could be safeguarded as “safety populations”. We estimate a change in IUCN Red List Category to LC.

*Tragelaphus derbianus*

Without knowing the population trend in the introduced population (Cuba), it is not possible to estimate if and of how much the IUCN RL assessment would change (IUCN SSC Antelope Specialist Group 2017c). We do not estimate a change in IUCN Red List Category.

*Petrogale penicillata*

Introduced only on two very small islands (Hawaii and New Zealand; Woinarski & Burbidge 2016c, Biancolini et al. 2021). Likely irrelevant. We do not estimate a change in IUCN Red List Category.

*Thylogale browni*

The introduced population is very close geographically to the native population (Leary et al. 2016b). Probably if the species is declining due to hunting on the mainland (Papua New Guinea), it will also be declining in the introduced population. However, nothing is known about EOO/AOO and population numbers, so we cannot assume that the rate of population decline is different between the native and alien populations. We do not estimate a change in IUCN Red List Category.

*Thylogale brunii*

Introduced only on one very small island (Kai Kecil Island; Indonesia; Leary et al. 2016). Likely irrelevant. We do not estimate a change in IUCN Red List Category.

*Phascolarctos cinereus*

The species has been introduced to various Australian islands and mainland Australia, and some populations are invasive and thus likely to have stable or positive population trends (Woinarski & Burbidge 2020). But it is not clear from the assessment whether these populations are included or not. Moreover, in the "In-place species management" section of the IUCN RL assessment, it is marked "Successfully reintroduced or introduced benignly: Yes" (Woinarzi & Burbridge 2020). We do not estimate a change in IUCN Red List Category.

*Lepus corsicanus*

In the main part of the introduced range (Corse Island; France) the species appears to be increasing, but detailed information on population trends or threats is missing (Randi & Riga 2019). We do not estimate a change in IUCN Red List Category.

*Eulemur albifrons*

Introduced to Nosy Mangabe (Madagascar) for conservation purposes according to DAMA (Biancolini et al. 2021) and Red List assessment (Borgerson et al. 2020), but in the Red List map it is shown as native. It seems anyways that the alien population is included in the assessment (Borgerson et al. 2020). However, the island is likely too small to make a difference. We do not estimate a change in IUCN Red List Category.

*Eulemur fulvus*

Possibly the introduced one (Mayotte Island, France) is a hybrid population (Irwin & King 2020). We do not estimate a change in IUCN Red List Category.

*Macaca arctoides*

DAMA refers to an invasive population in Cuba in 2017 (Biancolini et al. 2021). In the IUCN RL assessment (dated 2020), the species is results to be introduced to Hong Kong (Chetry et al. 2020). Likely irrelevant. We do not estimate a change in IUCN Red List Category.

*Macaca leonina*

There are populations of *M. leonina* and *M. nemestrina* found on either side of the distribution limits in the Isthmus of Kra (Asia), but many of these populations are the result of release by humans (Boonratana et al. 2022). Likely irrelevant. We do not estimate a change in IUCN Red List Category.

*Trachypithecus auratus*

The introduced population on Lombok Island (Indonesia) is considered of uncertain origin in the IUCN RL assessment (Nijman 2021), but it is present in the geographic range part. The island is very small, and the introduction is likely irrelevant. We do not estimate a change in IUCN Red List Category.
